# Strain Matters: The 129S1/SvlmJ Mouse Model Reveals the Genetic and Inflammatory Susceptibility to Hypertensive Complications

**DOI:** 10.1101/2025.03.24.641145

**Authors:** Arthur Orieux, Romain Boulestreau, Marie-Lise Bats, Maxime Michot, Alexandre Boyer, Virginie Dinet, Juliette Vaurs, Pascale Dufourcq, Claire Peghaire, Cécile Duplaa, Thierry Couffinhal, Sébastien Rubin

## Abstract

**Background:** Hypertension is a leading cause of microvascular injury, yet the genetic determinants of organ-specific vulnerability remain poorly understood. Yest, we need good mouse models to investigate the complication of hypertension. This study investigates the role of genetic background in shaping hypertensive complications by comparing two mouse strains with divergent inflammatory responses.

**Methods:** Three-month-old 129S1/SvlmJ and C57BL/6J mice received 600 ng/kg/min of angiotensin II (AngII) or saline. We compared the consequences of ANG2-induced blood pressure elevation on kidney function, BBB intergrity and cardiac hypertropy. Blood pressure (BP) was assessed by telemetry. Vascular injury markers in the brain, heart, kidneys, and retinas were systematically evaluated.

**Results:** Both strains developed similar moderate hypertension with AngII. Only 129S1/SvlmJ mice exhibited spatial learning and memory deficits, blood-brain barrier hyperpermeability, astrocyte activation, retinal artery damage, hypertrophic cardiomyopathy, and renal podocyte lesions with urinary albumin/creatinine ratio (UACR) after AngII treatment. Transcriptomic analysis of brain microvessels highlighted strain-specific differences in gene regulation, particularly in inflammatory pathways, which may explain the higher vulnerability of 129S1/SvlmJ mice to hypertensive organ damage. These findings were supported *in vivo* by increased resident and perivascular macrophage recruitment in the brain of C57BL6/J mice under AngII compared to the 129S1/SvlmJ strain.

**Conclusion:** Our findings highlight the critical role of genetic background in shaping hypertensive complications. The 129S1/SvlmJ strain serves as a valuable model for dissecting the molecular mechanisms of hypertensive organ damage, emphasizing neurovascular inflammation as a potential therapeutic target.

**Translational Perspective:** This study highlights the 129/Sv mouse strain as a superior translational model compared to the widely used C57BL/6J strain, which, despite being a standard in cardiovascular research, fails to reliably reproduce severe hypertensive organ complications. The 129/Sv strain closely mimics human hypertensive damage, including cerebral small vessel disease, nephropathy, cardiomyopathy, and retinopathy. Transcriptomic analysis of cerebral microvessels identifies maladaptive inflammation as a critical mechanistic driver of susceptibility. These findings underline the clinical relevance of genetic predisposition, improving risk stratification and providing a robust preclinical platform to develop targeted anti-inflammatory therapies aimed at preventing hypertension-induced end-organ damage in patients.

## I. Introduction

Hypertension affects over 1.5 billion individuals globally and remains the leading modifiable risk factor for cardiovascular and renal diseases^1^. It is also the leading chronic risk factor for years of life lost, accounting for roughly 200 million years lost per year^2,3^. This leads to severe complications such as cerebral small vessel disease (cSVD), cognitive impairment, chronic kidney disease (CKD), and progressive renal failure. Indeed, even with optimal blood pressure control, many patients continue to develop severe complications, while others remain largely unaffected, suggesting important mechanisms independent of the mechanical effects of hypertension. The precise factors determining organ-specific susceptibility and resilience to hypertensive damage, particularly in the brain and kidneys, are poorly understood. Understanding these determinants is therefore critical to developing targeted therapeutic interventions that address the residual risk of hypertensive complications beyond mere blood pressure management^4^.

In the pursuit of identifying the mechanistic determinants of organ-specific vulnerability, Genome-wide association studies (GWAS) in humans have identified genetic polymorphisms associated with hypertension and its complications^5^. Yet, translating these findings into mechanistic insights has been challenging. GWAS provide limited causal understanding, and clinical studies often cannot manipulate specific pathways experimentally^6^. Consequently, preclinical models are essential for exploring the biological processes underlying hypertensive complications. While mouse models remain invaluable tools in hypertension research, existing models often fail to replicate the complexity of human hypertension and its diverse organ-specific damage^7^. Furthermore, the role of genetic background in driving differences in hypertensive target organ injury remains poorly understood.

To address this gap, we need good mouse models to investigate the complication of hypertension. We examined how genetic background affects hypertensive complications using a model of moderate hypertension induced by angiotensin II (AngII). By comparing two genetically distinct mouse strains—129S1/SvlmJ (129/Sv) and C57BL/6J—we observed striking differences in the severity and patterns of organ-specific damage. Notably, 129/Sv mice exhibited extensive damage to the brain, kidneys, retina, and heart, reflecting severe hypertensive complications in humans. In contrast, C57BL/6J mice appeared to be protected from such damage. Interestingly, the C57BL/6J strain, despite being commonly recommended and extensively used for cardiovascular research, may therefore be suboptimal for modeling hypertensive organ complications due to its inherent resistance to developing significant end-organ damage under hypertensive conditions.

Transcriptomic analyses identified inflammatory pathways as central mediators of these strain-specific vulnerabilities, further confirmed by direct tissue analysis, highlighting potential therapeutic targets.

## II. Methods

### Animals

129S1/SvlmJ (129/Sv) mice were obtained from *Charles River Laboratories*, strain code 287, while C57BL/6J mice were sourced from *The Jackson Laboratory,* strain code 000664. The mice are maintained on a standard laboratory diet (SAFE A04, Safe Laboratory, France) and are housed in a specific pathogen-free environment.

The animals were housed in a conventional facility on a 12-hour (light/dark) cycle, with water and food provided ad libitum. This study was conducted in accordance with the local Animal Care and Use Committee at Bordeaux University (Committee CEEA50) and the regulations in effect within the European Community for experimental animal use (L358-86/609/EEC). General surgical procedures were performed on mice (ages 3–12 months, weights 25–35 g) following the ARRIVE guidelines (https://www.nc3rs.org.uk/arrive-guidelines).

### AngII-induced hypertension

Moderate hypertension was induced in three-month-old male mice by subcutaneous infusion of angiotensin II (AngII, Sigma-Aldrich, USA; A9525-50MG) using osmotic pumps (ALZET 1004, Alzet, USA). The pumps were preloaded to deliver AngII at a rate of 600 ng/kg/min for 28 days (D28). Control mice (SHAM) received an equivalent saline infusion (0.9%). The day of pump implantation was designated as day 0 (D0). Mice were anesthetized with 1.5% isoflurane in oxygen during pump implantation, and perioperative analgesia was provided through subcutaneous buprenorphine (0.05 mg/kg) (Ceva Santé Animale, Libourne, France). The pumps were implanted subcutaneously in the dorsal region to minimize handling stress. Mice were monitored postoperatively for recovery and weight stability.

### Blood pressure measurement

A catheter (PA-C10, AD Instrument, New Zealand) is inserted into the carotid artery, and the transmitter was placed subcutaneously. After five days, mice were placed onto RPC-1 Single Receiver (AD Instrument, New Zealand). Data were acquired every week over one hour during the day using LabChart software (v7, AD Instruments, New Zealand) to calculate mean systolic blood pressure (sBP).

### Brain damage assessment

#### Behavioral test

The Morris Water Maze (MWM) test was conducted from day 21 to 28 after hypertension onset, as previously described8, to assess spatial learning and memory. Before testing, visual acuity was confirmed using the visual cliff test. Briefly, mice underwent four acquisition trials daily over four consecutive days, learning to locate a submerged platform. Latency to reach the platform was recorded, with shorter latencies indicating better performance. Three days later, memory retention was evaluated by a 90-second Probe Test without the platform, measuring mean proximity to the original platform location. All behavioral data were collected and analyzed using EthovisionXT v16 software (Noldus Information Technology, Wageningen, Netherlands)^8,9^.

#### Cerebral blood flow

Laser-Doppler imaging (MOORLDI2-IR™, Moor Instruments, UK) was used to measure cerebral blood flow (CBF) at baseline and after 28 days of infusion. Mice were anesthetized during measurements, and body temperature was maintained on a heating platform.

#### Blood-brain barrier permeability assessment *in vivo*

As previously described^10^, mice were anesthetized, and 75 µl of TMR dextran 3 kDa lysine-fixable tracer (Texas Red™, 3000 MW, Lysine Fixable, ThermoFisher Scientific, Cat# D3328, MA, USA) was injected intravenously (retro-orbital). Twenty minutes later, a 300 μL blood sample was collected via cardiac puncture and centrifuged at 10,000 x g for 10 minutes at 4 °C to prepare the serum. The mice were euthanized immediately after collecting the blood sample. One hemibrain, devoid of olfactory lobes and cerebellum, was collected, and its weight was recorded before being snap-frozen in liquid nitrogen. Tissues were homogenized in cold 1X phosphate-buffered saline (PBS) using a Tissue Lyser (Qiagen, Germany) and centrifuged at 15,000 g for 20 minutes at 4 °C. The supernatant was then collected. Fluorescence measurement (RFUs) of 50 μL of diluted serum and brain supernatant was performed using a Spark^®^ Multimode Microplate Reader (Tecan). We calculated a permeability index (PI) expressed in mL/g (tissue RFUs/g tissue weight) / (serum RFUs/mL serum) for each animal. The PI of each animal in the groups was divided by the mean PI of the SHAM group, setting the SHAM group mean to 1 while maintaining the inter-animal variation within the control group. This transformation yields values for the AngII group relative to the SHAM group, which is set to 1 for each strain.

#### Comprehensive three-dimensional analysis of brain (CUBIC)

Mice were transcardially perfused with a wash solution (0.9% NaCl + heparin 20 U/mL) and then fixed using 4% PFA. Post-perfusion, the brains were harvested, and 400 µm-thick coronal sections of the parietal area were prepared using a vibratome.

The brain slices were subjected to tissue clearing and immunolabeling using the CUBIC method^11^. Briefly, the sections were incubated in progressive CUBIC-1 reagent concentration for 36 hours at 37°C to facilitate tissue clearing. After clearing, the slices were incubated with the primary antibody for 48 hours at 4°C with gentle shaking at 37°C. Then, brain slices were incubated with primary antibodies: GFAP (1:400, ThermoFisher Scientific, 130300, MA, USA), PODOCALYXIN (1:400, R&D Systems, AF1556, Minneapolis, MN, USA), IBA1 (1:400, Fujifilm Wako, 019-19741, Osaka, Japan), CD206 (1:400, R&D Systems, AF2535, Minneapolis, MN, USA). The slices were washed and incubated with the conjugated secondary antibodies (Alexa Fluor, ThermoFisher Scientific, MA, USA) under the same conditions for 48 hours. Finally, the slices underwent a second clearing step using progressive CUBIC-2 reagent concentration for 36 hours at room temperature. The cleared sections were imaged via confocal microscopy for 3D reconstruction of labeled structures.

Quantifications of astrocyte activation, perivascular macrophages, and resident macrophages were carried out using Fiji/Image^12^. Brain slices from five mice at each time point (SHAM and D28) were imaged, with five tiles in the parietal region analyzed per mouse. Images were acquired under consistent parameters, including laser power, exposure, pinhole size, and image dimensions. A Z projection was applied to 3D stacks for 2D representation. An automatic threshold was employed to segment activated astrocytes and resident macrophages from the background, using identical settings across all samples. For each mouse, the mean value of the five tiles was used to quantify the areas of astrocyte activation (GFAP staining) and resident macrophages (IBA1 staining). Perivascular macrophages coverage wasquantified by two blind readers, who divided the number of labeled macrophages by the vessel length (μm) according to CD206 staining.

### Heart assessment

#### Echocardiography

Left ventricular (LV) ejection fraction and dimensions were measured on a high-resolution echocardiography (VEVO 2100, VisualSonics Inc., Toronto, Canada) with a 30 MHz transducer^13^. Mice were anesthetized using 1,5% oxygenated isoflurane by inhalation and were anchored to a warming platform in a supine position and prepared to minimize ultrasound attenuation. Uni’gel ECG (Asept Inmed, France) was applied to the thorax for optimal imaging. The ejection fraction was evaluated by planimetry as recommended^14^. Two-dimensional, long- and short-axis views were acquired to determine end-systolic (ESV) and end-diastolic (EDV) volumes using a cylindrical-hemi ellipsoid model. LV ejection fraction was calculated as: (EDV-ESV)/EDV x 100. The cardiac wall thickness (left ventricular posterior wall (LVPW), interventricular septum (IVS), and left ventricular internal diameter (LVID) were calculated by tracing wall limits in both the long and short axis views. LV-mass was corrected for body weight and expressed as LV-mass-index (LVMass) (mg/g)^15^.

### Kidney assessment

#### Blood and urinary tests

Blood samples were collected in heparinized tubes. After centrifugation (2,500 x g for 10 minutes), plasma creatinine and blood urea nitrogen (BUN) were measured using an enzymatic method (Architect C16000, Abbott Diagnostics, Rungis, France). Urinary creatinine (UCr) was measured using the same enzymatic method. Albuminuria was evaluated using the Mouse Albumin ELISA Kit (Crystal Chem, IL, USA; Cat# 80970), following the manufacturer’s instructions.

#### Electron microscopy

Animals were anesthetized. An anterior thoracoabdominal incision was performed, and the heart, brain, and kidneys were immediately immersed in cold 1X PBS, stopping the heartbeat. The left ventricle, the brain, and the kidney were then carefully sliced under an optical magnifier and fixed in 4% PFA/2% glutaraldehyde (#16220, Euromedex, France) at 4°C. Kidney sections were post-fixed at room temperature (RT) in 1% osmium tetroxide in the dark and dehydrated with a graded ethanol series. A final dehydration step is performed with acetone and embedded in Epon. Ultrafine sections of 70 nm thick were then collected on copper grids (Electron Microscopy Sciences, Hatfield, PA, USA) and observed with a transmission electron microscope Hitachi H7650 (Hitachi High Technologies, Tokyo, Japan).

#### qPCR and podocin staining

Gene expression analysis was performed on kidney samples (6 mice per group) from 129/Sv mice under SHAM and AngII conditions at day 28 using RT-qPCR. Total RNA from kidney mouse tissue was isolated by using the Direct-zol™ RNA MicroPrep kit (R2062, Zymo Research) and reverse transcribed into cDNA using the M-MLV Reverse Transcriptase (Promega). Quantitative real-time PCR was performed using the GoTaq qPCR master mix (Promega) on a QuantStudio™ 3 qPCR System (Thermo Fisher Scientific, MA, USA). Expression levels of key genes involved in podocyte and endothelial functions (*Nphs1*, *Nphs2*, *Thsd7a*, *NPR3*, *Wt1*, and *Flt1*) were normalized to cyclophilin A and compared between conditions (Supplementary data, Table S1).

Immunofluorescence staining was conducted to visualize glomerular structures in C57BL6/J and 129/Sv mice under SHAM and AngII conditions. Podocin (NPHS2) staining was performed using the anti-Podocin polyclonal antibody (PA5-79757, Thermo Fisher Scientific, MA, USA).

### Retinal assessment

#### Retinal vasculatures analysis

Mice were intravenously injected with 75 µl of dextran (D3328, Texas Red™, 3000 MW, Lysine Fixable, ThermoFisher Scientific, MA, USA). Twenty minutes later, eyes were collected and fixed in 4% PFA for 30 minutes. Immunostaining on the whole-mount retina was conducted as previously described^16^. Retinas were stained with conjugated isolectin-B4-FITC (L2895, Sigma-Aldrich, MO, USA, dilution 1:200) diluted in PBS-1% Donkey serum (D9663-10ML, Sigma Aldrich, MO, USA) for 2 hours at room temperature before being flat mounted with Fluoromount G mounting medium (00-4958-02, ThermoFisher Scientific, MA, USA). Retinal vasculatures were imaged and analyzed with a fluorescent Axio Observer microscope (Zeiss, Germany).

#### Blood-retina barrier analysis

Retinas were promptly removed just after death, fixed overnight in 4% PFA – 1X PBS solution at 4 °C, and cryoprotected in successive sucrose solutions (10%, 20%, and 30%) diluted in 1X PBS before being stored at −80 °C. Transverse retina sections (14 µm thickness) were cut using a Microm HM550 cryostat (Microm, Walldorf, Germany) at −20 °C. The sections were mounted onto Polysine^®^ glass slides (Polysine^®^ Adhesion slides, Thermo Scientific, MA, USA) and stored at −80 °C until further processing. For staining, nonspecific binding sites were blocked with 10% normal horse serum and 0.5% Triton X-100 in PBS for 1 hour at room temperature (RT). The sections were incubated with primary antibodies overnight (CRALBP 1/500, sc-48354Santa Cruz, France; lectin DyLight649, L32472, ThermoFisher Scientific, MA, USA). After washing with 1X PBS (three times for 10 minutes at RT), sections were incubated for 1 hour at RT with the appropriate secondary antibodies (donkey anti-mouse) conjugated to Alexa Fluor 488 (diluted 1/500, Invitrogen, France). Following washing, sections were mounted with a fluorescent aqueous mounting medium containing DAPI. Images were acquired with a confocal microscope (LSM 700, Carl Zeiss) using a 20× objective at 1024 × 1024 pixels resolution. All immunostaining experiments were repeated three to five times with different animals.

#### Tissue Collection, Fixation, and Histological Analysis

Mice were anesthetized with ketamine (250 mg/kg, i.p.) and xylazine (20 mg/kg, i.p.). Transcardiac perfusion was performed by inserting a blunted butterfly needle into the left ventricle after cutting the right atrium. For heart immunohistochemical analysis, whole hearts were arrested in diastole using 1 mol/L potassium chloride. PBS was perfused at a rate of 2 mL/min, followed by 4% PFA. The heart, kidneys, and brain were excised and post-fixed in 4% PFA at 4°C overnight before being embedded in paraffin. Seven μm paraffin heart sections were stained with wheat germ agglutinin (WGA) and DAPI. The diameter of cardiomyocytes was measured in high-magnification (40×) images from five randomly selected areas in five mice, analyzing 50 cells per mouse (250 cells per group). Picrosirius Red staining involved immersing tissue sections in a 0.1% Sirius Red solution in saturated picric acid for one hour, followed by washing in distilled water, dehydrating in absolute alcohol, clearing in xylene, and coverslipping^17^. Sagittal hemibrain slices were cut every 100 μm and stained with Pearls’ Prussian blue to detect microbleeds larger than 50 µm^2^, and hemalum-eosin to identify ischemic areas^18^. Following standard histological protocols, the kidney sections were stained with Periodic Acid–Schiff (PAS) to highlight glomerular and tubular basement membranes.

#### RNA extraction

On day 7 and day 28 after pump insertion, mice were euthanized by cervical dislocation. As previously described^19^, microvessels isolation was performed using 6 mice per group. All procedures were carried out in a cold environment to minimize cell activation. Ipsilateral cortices were homogenized in MCDB131 medium (ThermoFisher Scientific, 10372019, MA, USA) containing 0.5% fatty acid-free BSA (Sigma Aldrich, 126609, MO, USA). The homogenate was centrifuged at 2,000 g for 5 minutes at 4°C. The resulting pellet was suspended in 15% dextran (∼70 kDa, MilliporeSigma, 31390, MA, USA) in PBS and centrifuged at 10,000 g for 15 minutes at 4°C. The microvessels pellet was resuspended in MCDB131 with 0.5% fatty acid-free BSA and centrifuged at 2,000 g for 10 minutes at 4°C. RNA was extracted from cerebral-enriched microvessels (6 mice per group) using QIAzol lysis reagent (3722047, Qiagen, Germany) and the Qiagen RNeasy Mini Kit (Qiagen, Germany) as previously described. RNA purity was determined for RNA sequencing by assaying 1,5 μl of total RNA prep on a NanoDrop 8000 spectrophotometer.

#### Bulk RNA-sequencing

Total RNA integrity was checked using an Agilent Technologies 2100 Bioanalyzer with an RNA Integrity Number value (PCRq’UB platform, Bordeaux University). RNA sequencing bundle with PolyA was performed by Genewiz (Azenta Life Sciences, South Plainfield, NJ, USA). RNA library preparation was prepared following the manufacturer’s recommendations (KAPA mRNA HyperPrep Kit from ROCHE). Final samples from the pooled library prep were sequenced on Nextseq 500 ILLUMINA, 2×150bp corresponding to 30 million paired end reads per sample. The quality of raw data was evaluated with FastQC. Genetic and isoform abundances were quantified with rsem 1.2.28, before normalization with edgeR Bioconductor package^20^. Finally, differential analysis was conducted using the glam framework likelihood ratio test from edgeR. Multiple hypothesis-adjusted p-values were calculated using the Benjamini—Hochberg procedure to control FDR. Mammalian Phenotype Ontology (MPO) enrichment analysis was performed to identify phenotypic associations of differentially expressed genes (DEGs) and provide a functional interpretation of transcriptional changes. By linking gene expression to defined phenotypic traits, MPO allows for a more biologically relevant analysis of disease mechanisms. Gene enrichment was conducted using the MouseMine platform (https://www.mousemine.org), enabling the systematic identification of overrepresented phenotypic terms associated with the observed transcriptional alterations.

#### Statistical Analysis

Data were expressed as mean and standard deviation (SD) for normally distributed data or median and interquartile range (IQR) for non-normally distributed data. Parametric (unpaired t-test) or non-parametric (Mann-Whitney U test) tests were selected based on the data distribution for comparisons between the two groups. For multiple group comparisons, statistical analyses were performed using one-way ANOVA or repeated measures ANOVA, depending on the data structure. A two-way ANOVA was conducted to evaluate the main effects and interactions between factors for experiments with two independent variables (group and time). Post-hoc comparisons, such as Bonferroni or Tukey’s tests, were applied to identify specific group differences following significant ANOVA results. All p-values were two-tailed, with statistical significance defined as p[<[0.05. The sample size, what each n represents, the statistical tests used, and the result are indicated in each figure legend. Statistical analyses were performed using RStudio Statistical Software (v. 2024.09.1+394; Posit Software, PBC, Boston, MA) for transcriptomic analysis and GraphPad Prism 10 (GraphPad Software, Inc) for other analyses.

## III. Results

### C57BL/6 and 129/Sv display the same blood pressure elevation upon AngII exposure

To induce hypertension, we chose angiotensin II (AngII) infusion, a widely validated experimental model known to reliably produce moderate and stable elevation in blood pressure, closely mimicking human hypertension, and allowing controlled investigation of associated organ-specific complications. 129/Sv and C57BL/6J mice were implanted with an osmotic pump delivering 600ng/kg/min of AngII (AngII group) or a saline solution (SHAM group) (Figure 1A). At baseline, sBP was 105.9 ± 7.9 in the C57BL/6J strain and 110.7 ± 8.9 mmHg in the 129/Sv strain (p = 0.32). Under AngII infusion, sBP rose significantly and remained stable in both strains at day 7, day 14, and day 21 (two-way ANOVA, p = 0.090 for weeks and p=0,14 for strains) (Figure 1B) with no significant difference at any time point (e.g., 141.5 ± 6 mmHg in C57BL/6J vs. 143.4 ± 11.8 mmHg in 129/Sv at day 7; p = 0.510). At day 28, all mice were alive and their weights were similar between the two groups (28.5 ± 3.4 g for C57BL/6J vs 28.3 ± 3.3 g for 129/Sv; p = 0.85).

**Figure 1.**
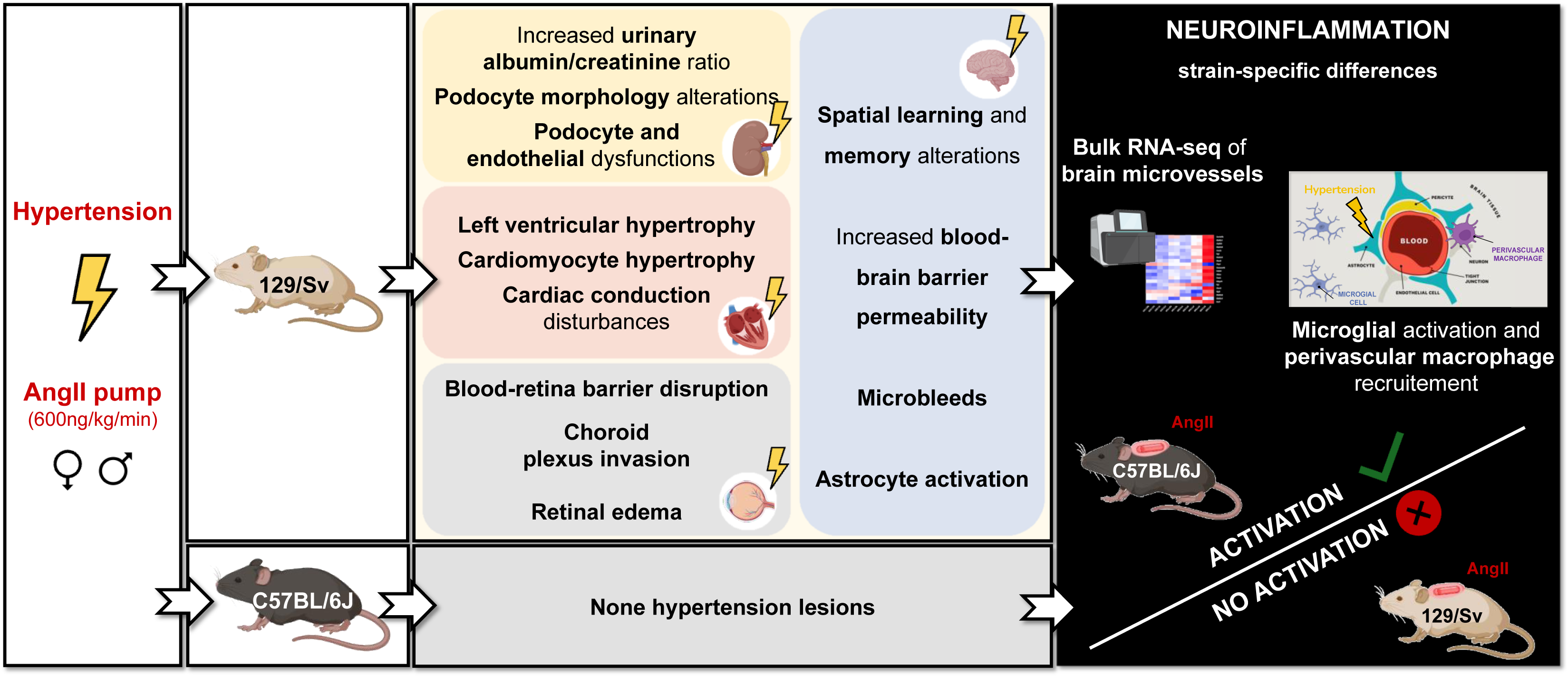
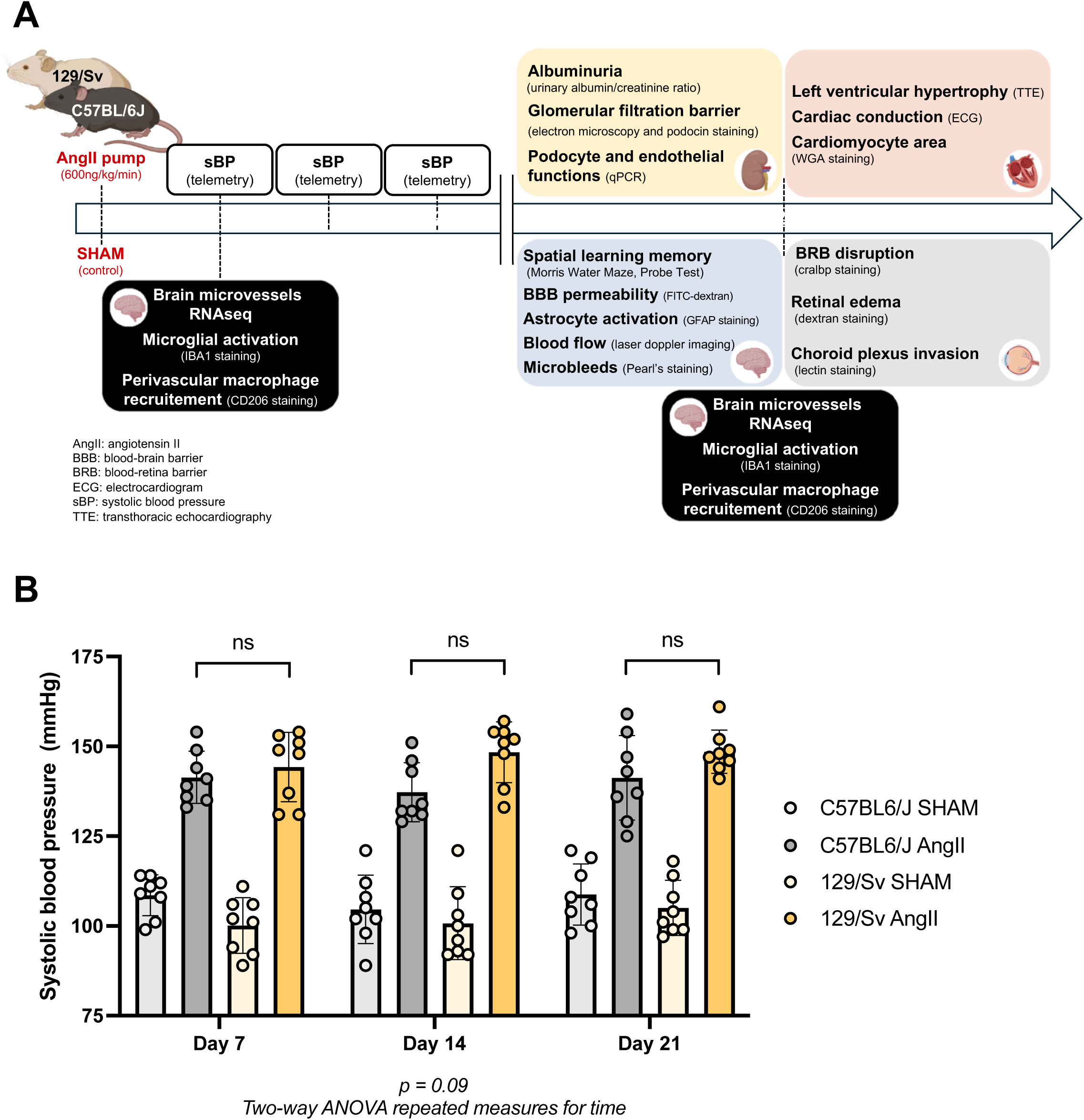
Experimental timeline and systolic blood pressure in 129/Sv and C57BL/6J mice after saline 0.9% (SHAM) or AngII infusion (A) Schematic representation of the experimental timeline. AngII (600 ng/kg/min) or saline (SHAM) was infused via osmotic minipumps from day 0 (D0) in 129/Sv and C57BL6/J strains. Systolic blood pressure (sBP) was measured weekly using invasive telemetry. At D28, the four target organs of hypertension (brain, heart, kidney, and retina) were evaluated through various analyses. Transcriptomic analyses were performed at D7 and D28. (B) sBP was measured with invasive telemetry at D7, D14, and D21 across four groups of mice (C57BL6/J SHAM, C57BL6/J with AngII, 129/Sv SHAM, and 129/Sv with AngII). Under AngII treatment, the 129/Sv and C57BL6/J mouse strains show increased sBP. However, as a two-way ANOVA (p = 0.09) indicates, this elevation does not significantly vary over time between the two strains.

### Only 129/Sv mice exhibit cognitive impairment, blood-brain barrier disruption, microbleeds, and astrocyte activation

In humans, hypertensive brain injury often manifests as cerebral small vessel disease (cSVD), characterized by cognitive impairment leading to vascular dementia. A central pathological feature of cSVD is disruption of the blood-brain barrier (BBB). We therefore examined cognitive functions, BBB integrity, and related cerebral lesions in hypertensive mice to assess strain-specific vulnerabilities and recapitulate human disease patterns.

In hypertensive C57BL/6J and 129/Sv mice, cognitive functions, particularly hippocampal memory integration, were tested using the Morris Water Maze starting at day 21. Visual acuity was assessed before the Morris Water Maze using the visual cliff test, confirming that no mice exhibited visual impairments. In the Morris Water Maze, no significant differences in latency or velocity were observed between SHAM and AngII-treated mice in the pre-training phase, indicating unaffected locomotor activity and learning performance (Supplementary data, Figure S1). After 21 days of AngII treatment, the 129/Sv AngII group demonstrated significant deficits in spatial learning and memory compared to 129/Sv SHAM, highlighting marked cognitive impairments in this strain. In the 129/sv strain, both groups performed similarly on day 1. Only 129/Sv SHAM mice showed improvement over time (p = 0.0008) (Figure 2A). Conversely, no significant cognitive impairment was observed in C57BL/6J AngII mice compared to SHAM, which showed similar performance in MWM (p = 0.18) (Figure 2B). In the Probe test (28 days after treatment (AngII or saline)), 129/Sv SHAM mice had significantly lower proximity measures than 129/Sv AngII (p = 0.047) (Supplementary data, Figure S2).

**Figure 2.**
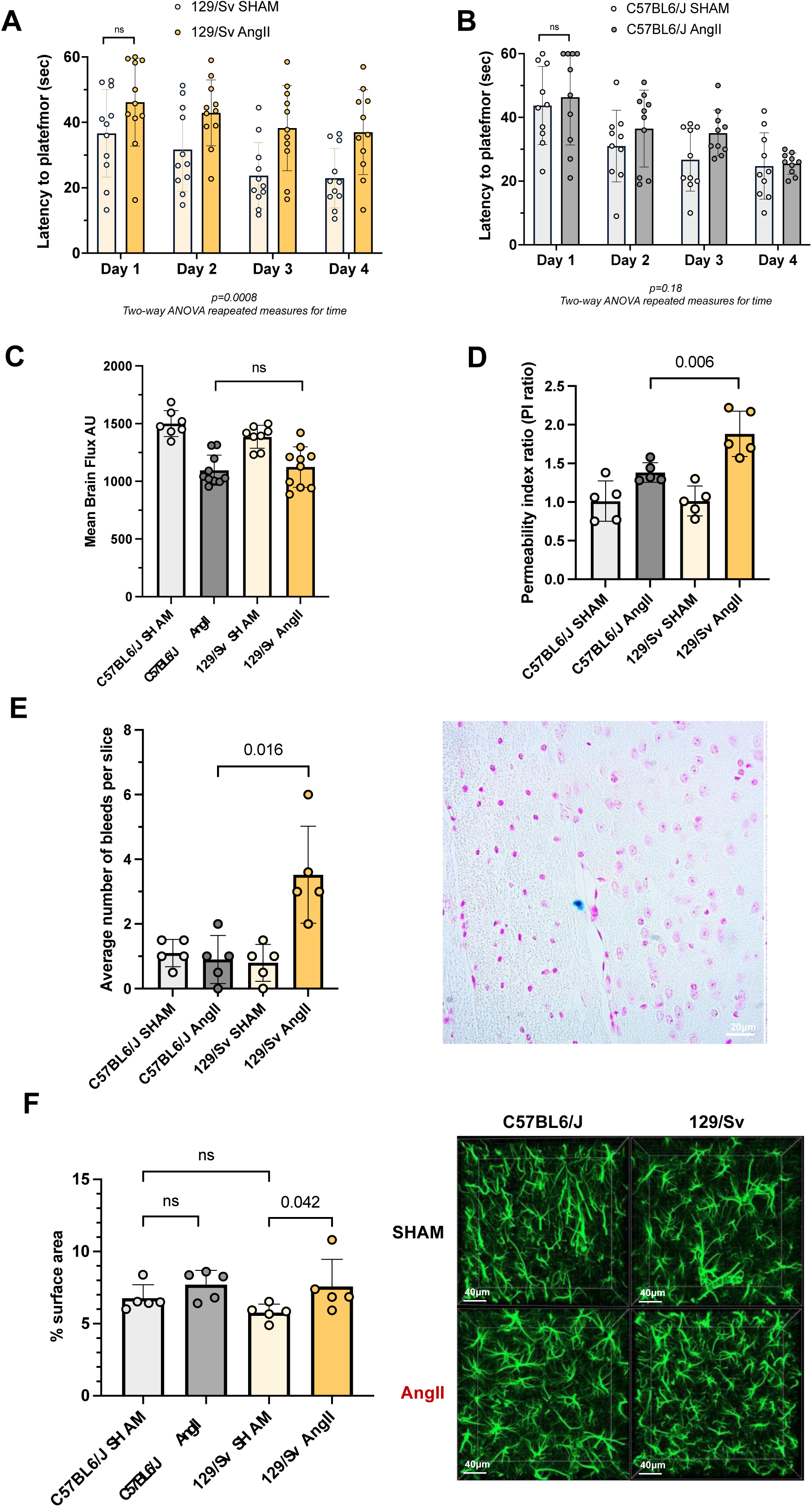
Brain effect of AngII infusion in both strains. (A) and (B) Morris Water Maze test showed similar spatial learning and memory in C57BL/6J mice under AngII compared to SHAM, but a disability in 129/Sv mice under AngII compared to their respective SHAMs. Graph shows the latency to reach the hidden platform (expressed in seconds) over 4 training days. Two-way ANOVA with repeated measures for time. (C) Mean brain CBF was similar in both AngII groups (1097 ± 131.6 AU in C57BL/6J vs 1124 ± 112.3 AU in 129/Sv; p = 0.70) (D) A significant increase in blood-brain barrier permeability was only found in 129/Sv mice after AngII infusion (permeability index ratio with 3kDa dextran) (E) Illustrative Pearl’s staining image of the brain from 129/Sv hypertensive mice. Using a systematic analysis of slices every 100 µm with Pearl’s staining, we found an increased average of microbleeds only in 129/Sv mice after AngII infusion (microbleeds larger than 50 µm^2^). (F) Representative immunofluorescence image of hippocampus area from both strains under two conditions. 129/Sv mice expose an astrocyte activation profile in hippocampal area (GFAP staining) after AngII infusion

CBF has been described as altered in vascular cognitive impairment^21^. We measured CBF before sacrifice (28 days of AngII). In both strains, AngII led to a reduction in brain CBF, but this decrease was similar between groups (1097 ± 131.6 AU in C57BL/6J vs 1124 ± 112.3 AU in 129/Sv; p = 0.700) (Figure 2C).

An increased brain-blood barrier (BBB) permeability was identified as one of the earlier BBB dysfunctions in the context of aggression^22^. We evaluated BBB permeability after 28 days of AngII or saline in both strains, using a 3kDa fluorescent dextran injected 20 minutes before the sacrifice. Results were expressed as the permeability index (PI) ratio of the AngII group relative to the SHAM group in each strain. After AngII infusion, the PI ratio was 1.33 ± 0.15 vs. 1.9 ± 0.30 (p = 0.006) in C57BL/6J vs 129/Sv, respectively (Figure 2D), showing a significant increase in BBB permeability in 129/Sv strain as compared to C57BL/6J. We did not find any vascular extravasation of larger molecules, including IgG (150 kDa) and fibrinogen (340 kDa), in immunostaining of brain paraffin slices (n = 5) (data not shown).

Cerebral lesions associated with hypertension are mostly small vessel lesions characterized by microinfarction and microbleeds^23^. Microbleeds were significantly more frequent in the 129/Sv AngII group than in the C57BL/6J AngII group (3 [2-4.75] vs. 1 [0.25-1.50]; p = 0.016) (Figure 2E). However, we did not find any ischemic areas in any group or strain (data not shown).

GFAP staining is a reliable marker for astrocyte activation, which can be induced by increased BBB permeability^24^. In the 129/Sv strain, a significant increase in GFAP staining was found after AngII infusion compared to C57BL/6J. In the hippocampal area, the mean GFAP surface area in the 129/Sv strain was 5.77 ± 0.59 % in SHAM condition and 7.47 ± 1.10 % after 28 days of AngII (p = 0.042) (Figure 2F).

### Only 129/Sv mice develop severe albuminuria and podocyte injury without nephroangiosclerosis

In humans, hypertensive kidney injury commonly manifests as hypertensive nephropathy, often characterized by low grade albuminuria. We therefore assessed renal function, albuminuria, and podocyte integrity in hypertensive mice to determine whether strain-specific susceptibilities mirror these clinically relevant patterns.

Renal function assessed by plasma creatinine dosage and blood urea nitrogen (BUN) was comparable in both strains in the SHAM group and remained unaffected in the AngII group.

In C57BL/6J mice, the urinary albumin/creatinine ratio (UACR) remained low and stable following AngII infusion (SHAM: 5.5 ± 8.1 mg/mmol; D28: 12.5 ± 5.3 mg/mmol). By contrast, 129/Sv mice exhibited a dramatic increase in UACR at D28 (214.7 ± 90.8 mg/mmol) compared to their SHAM counterparts (8.2 ± 4.8 mg/mmol; p = 0.0079) (Figure 3A).

**Figure 3.**
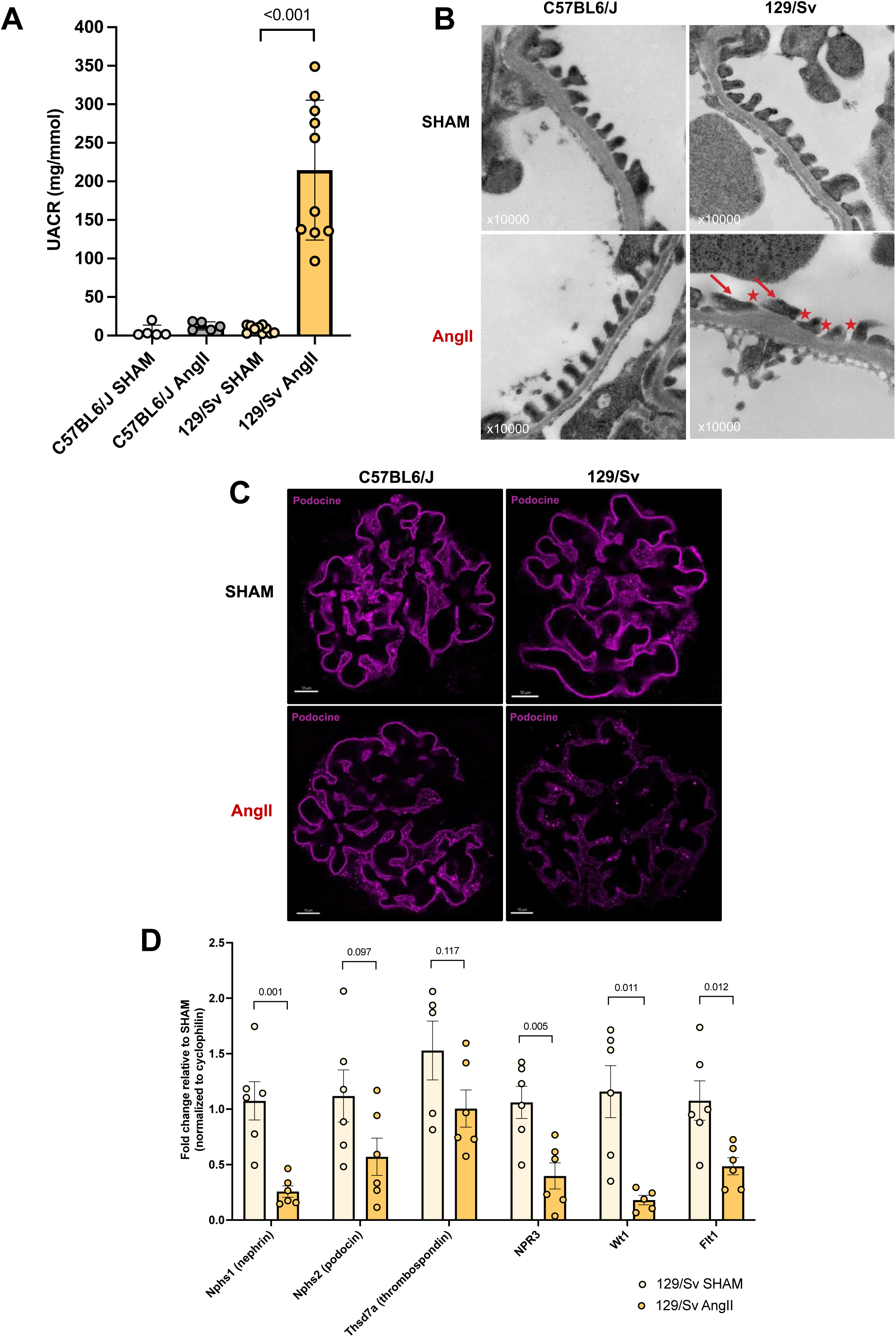
Kidney effect of AngII infusion in both strains. (A) 129/Sv mice exhibited a significant increase urinary albumin/creatinine ratio (UACR) at D28 (214.7 ± 90.7 mg/g) compared to the SHAM group (12.5 ± 5.3 mg/g, p < 0.001) (B) Representative electron microscopy image of the kidney from both strains under two conditions. After 28 days of AngII infusion, abnormalities in podocyte foot processes (red arrow) and the slit diaphragm (red star) are observed in 129/Sv mice compared to C57BL6/J. (C) Representative immunofluorescence image of a glomerulus from both strains under two conditions. Podocin staining: continuous linear staining in C57BL6/J with AngII vs. granular pattern staining in 129/Sv with AngII (D) qPCR of total kidney: decreased expression of podocyte and endothelial markers in 129/Sv strain after AngII. See Supplementary Table for the list of primers.

Given the previously described link between podocyte alteration and albuminuria^25^, we assessed podocyte morphology using electron microscopy. Notable abnormalities, including flattening of podocyte feet and changes in the slit diaphragm, were observed after 28 days of AngII infusion in 129/Sv mice but not in C57BL/6J mice (Figure 3B). Immunofluorescence staining using the anti-Podocin antibody revealed a disruption of the continuous linear pattern observed in SHAM mice, with a shift towards a more granular and fragmented distribution in 129/Sv AngII-treated mice, suggesting strain-specific differences in podocyte integrity and glomerular structure (Figure 3C).

RT-qPCR analysis revealed significant downregulation of Nphs1 (podocyte filtration), NPR3 (sodium regulation), Wt1 (podocyte development), and Flt1 (vascular permeability) in 129/Sv mice under AngII compared to SHAM (p < 0.05), confirming impaired podocyte and endothelial functions (Figure 3D).

Other vascular lesions, including ischemic glomerulus and fibrosis, were systematically searched using periodic acid Schiff staining. For either strain, no evidence of nephroangiosclerosis was identified in the AngII group (Supplementary data, Figure S3).

### Only 129/Sv mice exhibit hypertensive cardiopathy

The cardiac consequences of hypertension are dominated by left ventricular hypertrophy and fibrosis. To evaluate these effects, echocardiography was conducted on both mouse strains following the initiation of AngII or saline infusion. After 28 days of AngII treatment, the 129/Sv mice showed signs of hypertrophic cardiomyopathy, as evidenced by a significantly increased LVMass index (6.92 ± 0.96 mg/g vs. 5.17 ± 0.58 mg/g at baseline; p = 0.005), while no notable changes were found in the C57BL/6J strain (6.01 ± 0.65 mg/g vs. 6.15 ± 0.94 mg/g at baseline; p = 0.78). In the 129/Sv AngII group, there was an increase in systolic and diastolic left ventricular posterior wall (LVPW) and interventricular septal (IVS) thickness, accompanied by a decrease in left ventricular internal diameter (LVID). Conversely, the C57BL/6J strain did not exhibit significant differences in these parameters under AngII treatment (Figure 4A).

**Figure 4.**
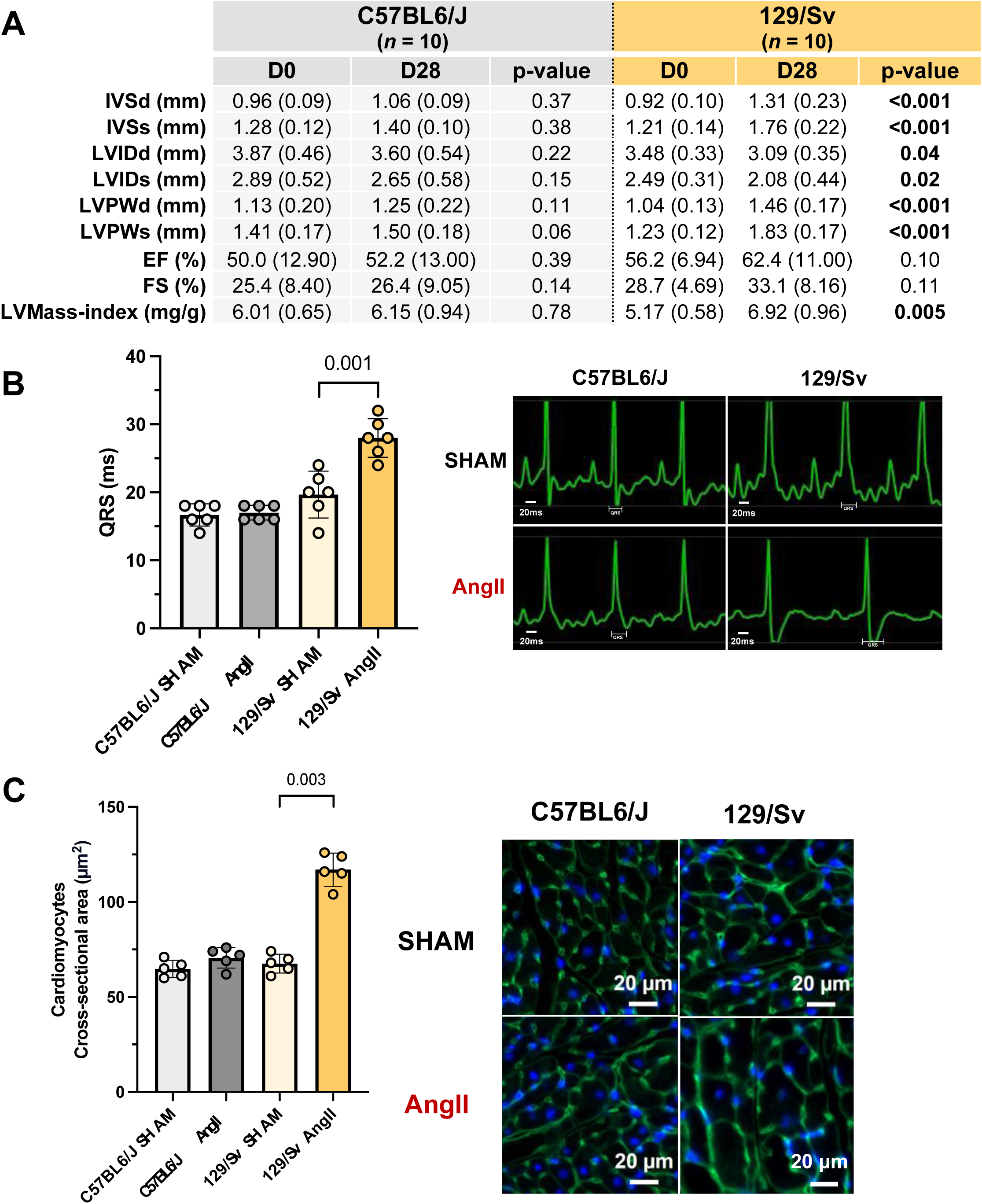
Heart effect of AngII infusion in both strains. (A) After 28 days under AngII, 129/Sv mice exhibited progressive hypertrophic cardiomyopathy contrary to the C57BL6/J strain. 129/Sv strain suffered from left ventricular hypertrophy with a corrected LV-mass index of 6.92 ± 0.96 mg/g vs. 5.17 ± 0.58 mg/g at baseline (p=0.005). (B) Illustrative electrocardiogram from both strains under two condition. Hypertensive 129/Sv mice exhibit cardiac conduction disturbances with prolonged QRS duration. (C) Representative immunofluorescence image of heart from both strains under two conditions. Significant increase in cardiomyocyte cross-sectional area (WGA staining) after 28 days of AngII infusion in 129/Sv vs. C57BL/6J strain (117.0 ± 8.7 vs. 67.6 ± 5.0 µm^2^; p=0.03). *EF: ejection fraction; FS: fractional shortening; IVSd: interventricular septal thickness in diastole; IVSs: interventricular septal thickness in systole; LVIDd: left ventricular internal diameter in diastole; LVIDs: left ventricular internal diameter in systole; LVMass-index: left ventricular mass index; LVPWd: left ventricular posterior wall thickness in diastole; LVPWs: left ventricular posterior wall thickness in systole*

In addition to structural changes, hypertensive 129/Sv mice exhibit cardiac conduction disturbances evidenced by a prolonged QRS duration (Figure 4B).

To further investigate myocardial remodeling, picrosirius red staining was used to evaluate cardiac fibrosis. Following AngII infusion, no fibrosis was detected in the cardiac tissue of either strain (data not shown).

Cardiomyocyte hypertrophy was evaluated using wheat germ agglutinin (WGA) FITC immunostaining. After 28 days of AngII treatment, 129/Sv mice had a significantly larger cardiomyocyte cross-sectional area than C57BL/6J mice (113.04 ± 17 vs. 72.22 ± 7 µm²; p = 0.03). However, no differences in cardiomyocyte cross-sectional area were noted between C57BL/6J and 129/Sv mice in the SHAM group (Figure 4C).

### Only 129/Sv mice develop blood-retinal barrier disruption

Hypertension can lead to various retinal complications, including hemorrhage, cotton wool spots, exudates, and papilledema in humans. At day 28, a disruption of the blood-retinal barrier was observed in hypertensive 129/Sv mice, as evidenced by cellular retinaldehyde-binding protein (CRALBP) immunostaining and choroid plexus invasion revealed by lectin staining. Notably, these changes were absent in hypertensive C57BL/6J mice (Figure 5A).

**Figure 5.**
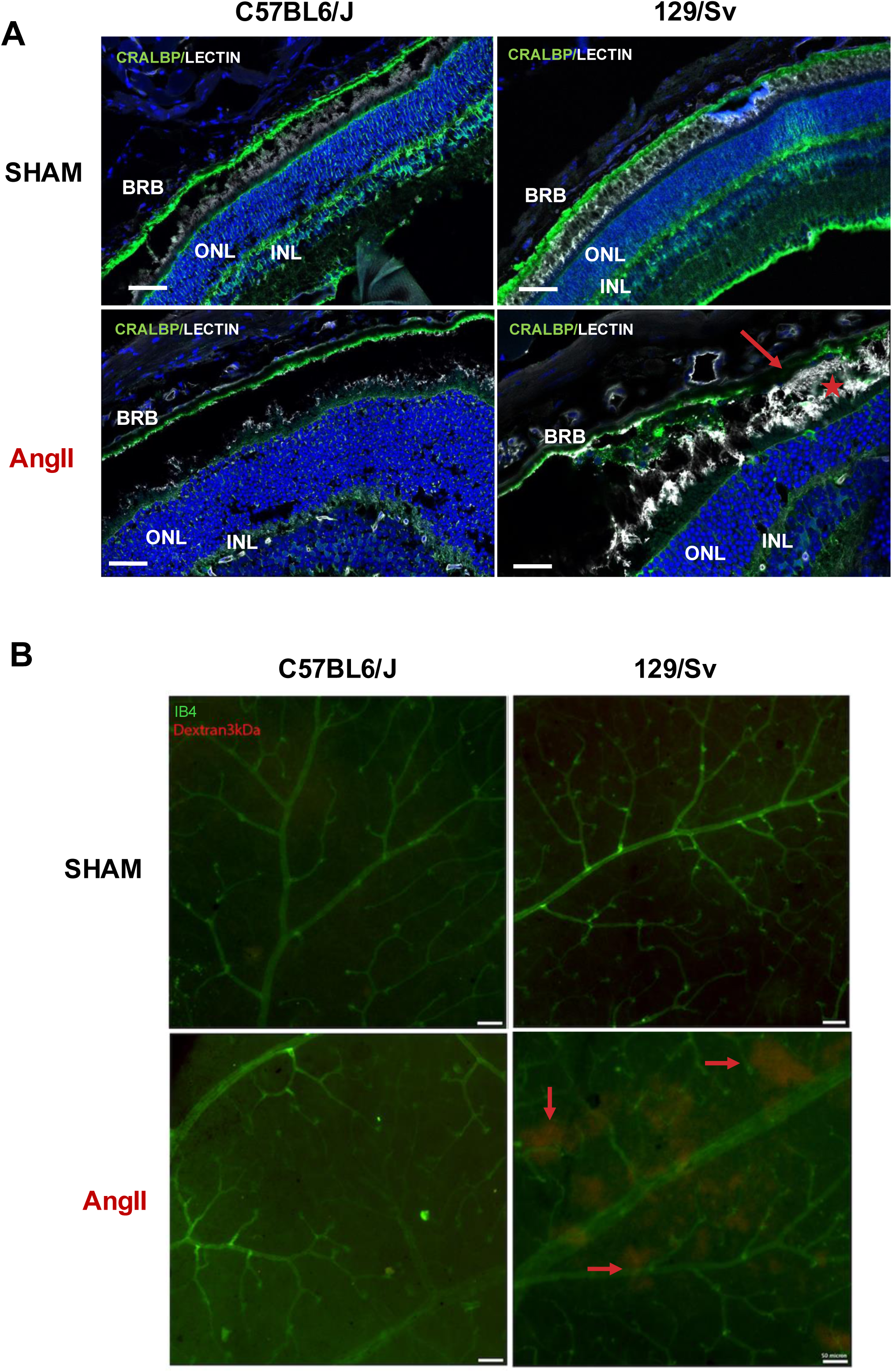
Retina effect of AngII infusion in both strains. (A) Representative immunofluorescence image of the retina from both strains under two conditions. Blood-retina barrier disruption (red arrow) was observed in 129/Sv mice with AngII-induced hypertension, along with choroid plexus invasion (red star), whereas no such disruption was seen in C57BL/6J mice with AngII. (B) After 28 days of AngII infusion, we found extravasation of dextran 3kDa (red arrow) on 129/Sv analyzed retinas (n=5) but none in C57BL6/J. *BRB: blood retina barrier; Ib4: Isolectin Ib4; INL: inner nuclear layer; ONL: outer nuclear layer*

We used TMR dextran three kDA lysine fixable as a tracer to further investigate retinal edema, a severe hallmark lesion. After 28 days of AngII treatment, we observed vascular extravasation of the dextran in all retinas analyzed from 129/Sv mice. In contrast, no extravasation was detected in the retinas of C57BL/6J mice (Figure 5B). The retinal lesions developed in 129/sv mice did not alter their visual acuity, measured by cliff test (data not shown).

### Female 129/Sv mice exhibit similar hypertensive organ damage as males upon 600 ng/kg/min AngII infusion

In the literature, a sexual dimorphism in the AngII-induced hypertension model is reported^26^. We decided to verify this hypothesis in a cohort of female mice. Similar to the findings in male mice, three-month-old 129/Sv female mice treated by AngII experienced organ injuries. In female 129/Sv mice, no cognitive tests were performed, and only tissue lesions were assessed. Histologic analysis revealed increased astrocytic activation under AngII (GFAP staining), with no significant differences compared to male 129/Sv mice in the SHAM condition. We observed elevated UACR with AngII treatment in female 129/Sv mice, with no significant differences compared to male mice of the same strain. Lastly, female 129/Sv mice developed hypertrophic cardiomyopathy under AngII infusion, as did male mice (Supplementary data, Figure S4).

### Transcriptomic analysis reveals strain-specific immune responses underlying susceptibility to hypertensive complications

To better understand why 129/Sv mice developed severe organ damage while C57BL/6J mice remained protected despite similar hypertension levels, we conducted transcriptomic analyses on brain-enriched microvessels isolated from both strains. We focused specifically on cerebral microvessels for this transcriptomic analysis because the brain exhibited the most striking and clinically relevant phenotype among the target organs, and cerebral microvascular injury represents a hallmark of hypertensive cerebrovascular disease in humans, providing a robust proof-of-concept model to understand strain-specific susceptibilities. Comparisons between the strains were performed at two critical time points: day 7, to capture early transcriptomic responses preceding the appearance of irreversible tissue lesions, and day 28, representing chronic stages of hypertension with established end-organ lesions. The volcano plots illustrate the differentially expressed genes (DEGs) between C57BL6/J and 129/Sv mice under AngII treatment at days 7 and 28 (Figure 6A).

**Figure 6.**
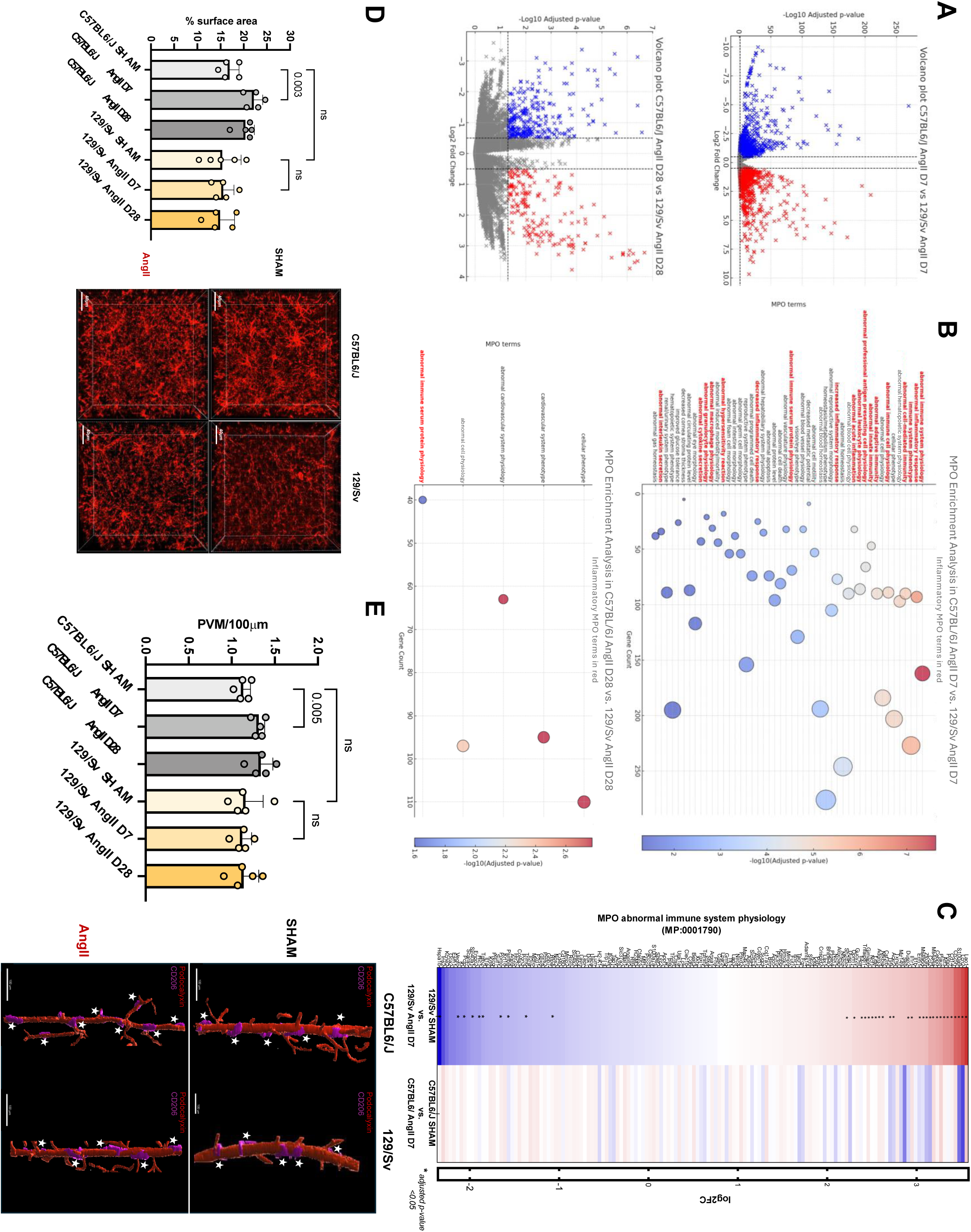
Transcriptomic and vascular differences between C57BL6/J and 129/Sv mice under AngII treatment (A) RNA-sequencing was performed on brain-enriched microvessel preparations from both strains under two conditions to investigate the biological processes underlying strain differences. Volcano plots illustrate the differentially expressed genes (DEGs), represented as log₂ fold change (FC) vs. –log₁₀ adjusted p-value, comparing C57BL6/J and 129/Sv mice under AngII at D7 (top) and D28 (bottom). Significant downregulated (log2FC <-1, in blue) and upregulated (log2FC >1, in red). In gray: non-significant genes. (B) Mammalian Phenotype Ontology (MPO) enrichment analysis of differentially expressed genes between C57BL6/J and 129/Sv mice under AngII at D7 (B) and D28 (C). Dot size represents gene count, and color indicates statistical significance (-log10 adjusted p-value). The MPO terms highlighted in bold and red font correspond to phenotypes related to inflammation. (C) Heatmap of differentially expressed genes between C57BL6/J and 129/Sv mice under AngII. The color scale represents log2 fold change, with red indicating upregulation (log2FC >1) and blue downregulation (log2FC <-1). Significant genes are marked with asterisks. (D) Representative immunofluorescence image showing microglial activation in the hippocampal area from both strains under two conditions. Quantification of IBA1-positive microglial activation in the hippocampal area. A significant increase in microglial activation is reported in C57BL/6J with AngII. (E) Representative immunofluorescence image showing perivascular macrophage coverage in the brain from both strains under two conditions. Quantification of CD206-positive perivascular macrophages in the parietal cortex. A significant increase in CD206-positive macrophages is observed in C57BL6/J mice with AngII.

To translate the observed gene-expression differences into meaningful biological insights, we performed a Mammalian Phenotype Ontology (MPO) enrichment analysis. The MPO database categorizes transcriptomic changes into known phenotypic consequences, facilitating interpretation of how differential gene expression could contribute mechanistically to observed organ-specific susceptibilities.

At day 7, comparison between strains revealed significant enrichment of immune-related and inflammatory MPO phenotypes, notably “abnormal immune system physiology” (MP:0001790, p < 0.0001) and “abnormal inflammatory response” (MP:0001845, p < 0.0001) specifically in the 129/Sv strain compared to C57BL/6J (Figure 6B). These results indicate that 129/Sv mice exhibit an early aberrant activation of inflammatory pathways, contrasting sharply with the controlled and protective low-grade inflammation observed in the C57BL/6J strain.

By day 28, strain-specific MPO differences were characterized by a shift towards cardiovascular, structural, and cellular remodeling phenotypes, reflecting progression from initial inflammatory responses to established structural vascular damage predominantly in the 129/Sv mice (Figure 6B). This temporal transition highlights the pivotal role of initial immune responses in determining subsequent susceptibility to chronic cerebrovascular damage.

Specifically, genes associated with the MPO term “abnormal immune system physiology” (MP:0001790) were significantly upregulated in 129/Sv mice treated with AngII at day 7 compared to C57BL/6J mice (Figure 6C), further confirming the maladaptive early inflammatory response occurring uniquely in 129/Sv mice. These findings collectively underscore how genetic background critically modulates the balance between protective and maladaptive inflammatory responses, thereby influencing susceptibility to hypertensive cerebrovascular complications.

### Only C57BL/6J mice exhibit protective microglial and perivascular macrophage recruitment in the brain

AngII was known to promote tissue inflammation cell recruitment^27^, and inflammation was described as an actor of hypertension pathogenesis and complications^28^. In line with the transcriptomic analysis, we wish to validate this difference *in vivo* in the brain. IBA1 is a marker of microglia (resident macrophages in the brain), and IBA1 staining was used to assess inflammatory cell recruitment in the brain^29^. Perivascular macrophages (PVM) expressing CD206 in the brain play a crucial role in immune surveillance and maintaining homeostasis within the central nervous system^30^.

After 28 days of AngII infusion, we found a significant increase in brain microglia staining using IBA1 staining in C57BL/6J in the hippocampal area compared with C57BL6/J SHAM mice (17.0 ± 2.0 % vs. 20.4 ± 1.9%; p=0.0263) (Figure 6D). Similarly, an increase in CD206 PVMs coverage was observed in the C57BL6/J strain under AngII with 1.13 ± 0.08 PVM/μm vs. 1.34 ± 0.13 PVM/μm in C56BL6/J SHAM mice (p=0.0227) (Figure 6E). On the contrary, no difference was observed in IBA1 and CD206 staining in the 129/Sv strain under AngII (vs. SHAM procedure).

## IV. Discussion

Hypertension is a major risk factor for cardiovascular and renal diseases, but the severity of organ damage varies widely among individuals. This variability suggests a strong genetic component influencing susceptibility to hypertensive complications. Our study emphasizes the critical influence of genetic background, showing that 129/Sv mice develop severe target organ damage despite similar increases in BP and low hypertension. In contrast, C57BL/6J mice remain largely protected.

Despite comparable hypertensive burden, 129/Sv mice exhibited widespread target organ damage, including cognitive impairment, BBB hyperpermeability, astrocyte activation, and increased microbleeds in the brain, along with retinal vascular permeability, hypertrophic cardiomyopathy, and podocyte injury-associated albuminuria in the kidneys. In contrast, C57BL/6J mice showed minimal or no such complications, reinforcing the hypothesis that genetic background dictates the severity of hypertension-related injury rather than the magnitude of BP elevation alone.

The immune response to AngII differs fundamentally between strains. In C57BL/6J, in response to low-dose AngII infusion, we observe a low-grade, controlled inflammation characterized by microglial activation and perivascular macrophage recruitment. This response appears adaptive and supports neurovascular homeostasis, as evidenced by preserved BBB integrity^31^.

Conversely, in 129S1/SvlmJ, there is a failure to mount a protective immune response early on, leading to BBB hyperpermeability, astrocytic overactivation, and cognitive impairment. This pattern closely resembles those described in cerebral small vessel disease (cSVD) in humans ^32,33^. In 129S1/SvlmJ, inflammation fails to act as an adaptive defense and instead appears dysregulated and maladaptive, driving progressive neurovascular damage. These results underscore the importance of inflammatory and immune-mediated mechanisms in modulating hypertensive injury and offer a dependable preclinical model for examining the genetic determinants of hypertension-induced end-organ damage.

BBB dysfunction has been reported as a delayed consequence of endothelial injury, possibly driven by AngII-mediated oxidative stress, aldosterone, endothelin-1 signaling, or recruitment of inflammatory cells^34,35^.

Glial cells play a fundamental role in maintaining neurovascular integrity and promoting neuroinflammatory tolerance^36^. In C57BL/6J mice, glial activation appears to be a compensatory mechanism that preserves neurovascular function, whereas in 129/Sv mice, the absence of this response contributes to greater susceptibility to hypertensive damage. Similar findings have been reported by Faraco *et al.*, who showed that AngII with salt-induced hypertension promotes perivascular macrophage recruitment and reactive oxygen species production, leading to BBB destabilization^7^. However, in C57BL/6J mice, this inflammatory response occurs without leading to BBB hyperpermeability, suggesting a protective rather than a detrimental effect. These findings highlight a fundamental distinction in how inflammatory pathways influence hypertensive outcomes across genetic backgrounds.

We analyzed cerebral microvessel transcriptomes to explore the mechanisms underlying these differences in susceptibility. By day 7 post-AngII, 129/Sv mice displayed a marked upregulation of immune-related pathways, particularly those linked to inflammation and endothelial dysfunction. In contrast, C57BL/6J mice exhibited a more controlled immune activation with fewer transcriptomic alterations. Uncontrolled inflammation in 129/Sv mice appears maladaptive, contrasting with the low-grade neuroinflammatory response in C57BL/6J, which may enhance vascular resilience. By day 28, this acute immune activation had transitioned towards a fibrotic and vascular remodeling phase, indicating that early inflammatory dysregulation in 129/Sv leads to long-term structural vascular damage. This subtle yet sustained activation in C57BL/6J could serve as a physiological protective mechanism, while the poorly regulated inflammation in 129/Sv may lead to maladaptive damage. A finely tuned inflammatory environment appears crucial in mitigating hypertensive injury.

In contrast, 129/Sv mice fail to mount a protective neuroinflammatory response, instead exhibiting a delayed, exaggerated, and maladaptive immune activation that exacerbates end-organ damage. The absence of protective neuroinflammatory recruitment in this strain may fail to counteract the detrimental effects of hypertension, further worsening end-organ damage. This aligns with prior studies highlighting the role of immune cell activation in determining susceptibility to hypertensive injury^37,38^. These findings suggest that a finely balanced inflammatory response acts as a safeguard against hypertensive damage, whereas dysregulated immune activation exacerbates cerebrovascular injury.

This pattern was consistently observed in male and female mice, challenging the hypothesis that female mice are inherently shielded from AngII-induced hypertension^26^. The consistency of these findings across sexes strengthens the reliability of this hypertensive model.

Our study identifies the 129S1/SvlmJ strain as a relevant model for severe hypertensive complications, mirroring human lesions in cSVD, cardiac hypertrophy, and renal dysfunction, further underscoring its translational relevance. Furthermore, our findings suggest that targeting inflammatory pathways could provide novel therapeutic strategies for mitigating hypertensive end-organ damage. Given the strain-specific differences observed in perivascular macrophage coverage and endothelial transcriptomic responses, interventions aimed at modulating neurovascular inflammation could be explored to enhance resilience in susceptible populations.

Finally, these findings reinforce the necessity of incorporating genetic background considerations into preclinical hypertension research. Many current hypertensive models rely predominantly on C57BL/6J mice, which, as demonstrated here, may underestimate the extent of end-organ damage observed in human pathology. Hamzaoui *et al.* reported that compared to C57BL/6J, the 129/Sv strain exhibits more severe cardiovascular injury following 5/6 nephrectomy, further supporting its utility in modeling hypertensive organ damage^10^.

These findings provide a unique platform for dissecting the interplay between genetics and hypertension-induced organ damage. Moreover, they position the 129S1/SvlmJ strain as a critical model for studying hypertensive complications and developing targeted therapeutic interventions.

In conclusion, this study demonstrates that genetic background significantly influences the severity of hypertensive complications, with 129/Sv mice showing widespread end-organ damage compared to the relatively protected C57BL/6J strain. Our transcriptomic analysis highlights strains’ differing inflammatory and endothelial responses, providing a mechanistic basis for these disparities. By identifying 129/Sv as a highly susceptible model, we establish a robust preclinical framework to uncover the pathophysiological drivers of hypertension-induced end-organ damage and to accelerate the development of targeted therapeutic interventions.

## Supporting information

qPCR amorces

## Acknowledgments

We sincerely thank Philippe ALZIEU, Sylvain GROLLEAU, Maxime DAVID for their technical assistance and animal care; Christelle BOULLE and Hélène AOUIZERATE for administrative assistance.

We thank Prof. Céline GUILBEAU-FRUGIER (UMR 1297, Toulouse, France) for Electron Microscopy images

This work benefited from equipment and services from the PCRq’UB platform (Bordeaux University).

## Sources of Funding

Arthur ORIEUX (salary grant): MD-PhD student grant – CHU de Bordeaux et Université de Bordeaux

Bourse Société Francophone de Néphrologie Dialyse et Transplantation (SFNDT) 2023

This project is supported by a grant overseen by the French National Research Agency (ANR) as part of the Investment for the Future Programme ANR-18-RHUS-0002 (SHIVA Project) and by the Precision and Global Vascular Brain Health Institute (VBHI) funded by the France 2030 IHU3 initiative

## Supplemental Material

Figure S1-S4

Table S1

Forward and reverse primer sequences for genes used in qPCR

---

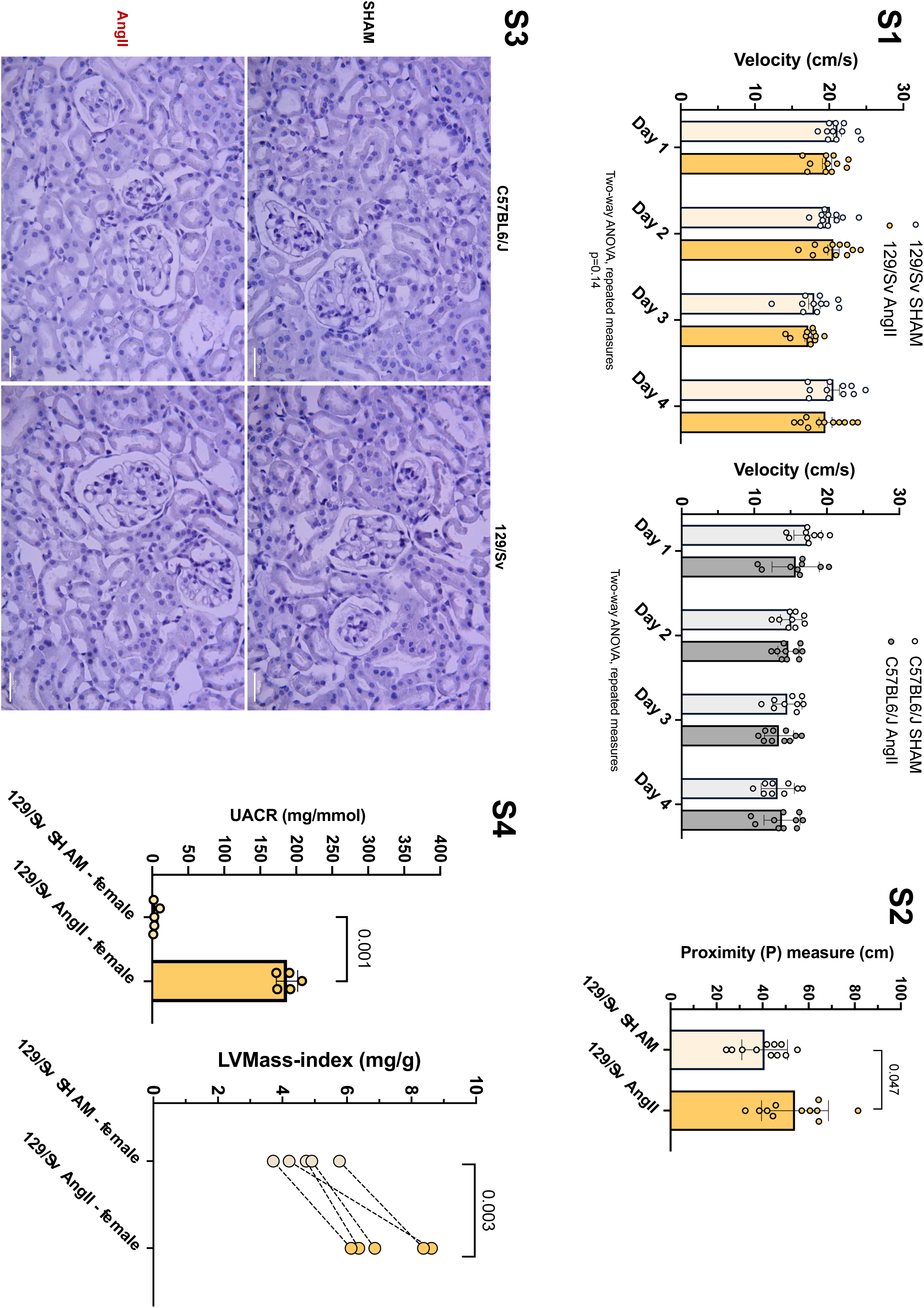

