## Supplementary material for "Strain Matters: The 129S1/SvlmJ Mouse Model Reveals the Genetic and Inflammatory Susceptibility to Hypertensive Complications": qPCR amorces

**Forward and reverse primer sequences for genes used in qPCR**

| **Gene** | **Forward** | **Reverse** |
| --- | --- | --- |
| **Cyclophilin A** | AGCTAGACTTGAAGGGGAATG | ATTTCTTTTGACTTGCGGGC |
| **Nphs1 (nephrin)** | TGGCGATTCCTGCCTCCGTT | TTCTGCTGGGAGCCCTCGTT |
| **Nphs2 (podocin)** | GTGTCCAAAGCCATCCAGTT | GACCTTTCCTTCTCGTAACG |
| **Thsd7a** | ACCTGGGTTTATGGTGTCG | GGTTTGAGGCGGTTGTT |
| **NPR3** | GTACTCAGAGCTGGCTACAGCA | CGTTGGCATCTATGGACACCTG |
| **Wt1** | AGTGAAATGGACAGAAGGGCAGA | TCCAGATACACGCCGCACAT |
| **Flt1** | CCACCTCTCTATCCGCTGG | ACCAATGTGCTAACCGTCTTATT |
